# Mechanisms of Interaction of Staphylococcus Aureus With Human Mesenchymal Stem Cells and Their Differentiated Phenotypes

**DOI:** 10.1101/2020.01.09.900373

**Authors:** Nibras Khamees, Darryl J. Hill, Wael Kafienah

## Abstract

Mesenchymal stem cells (MSCs) are multipotent cells commonly derived from the bone marrow, adipose tissue and placenta. Human bone marrow derived MSCs migrate to a site of injury, release proinflammatory cytokines and modulate T-cell proliferation. At sites of injury, MSCs may well encounter bacterial pathogens most commonly the Gram positive pathogen *Staphylococcus aureus*. However, the precise molecular mechanism(s) of this interaction remain to be elucidated. In the present study we aim to show if a direct interaction occurs between *S. aureus* and bone marrow derived MSCs and identify if MSCRAMMs have a role in this interaction. We further aim to compare *S. aureus* interaction with cells that differentiate from MSCs, namely; osteoblasts, adipocytes and chondrocytes, since MSCs co-exist in the niche of these cells. Our results showed that S. aureus is able to interact with MSCs in the form of adhesion and invasion to the cells, and that this interaction is largely dependent on the expression of fibronecting-binding protein (FnBP) by S. aureus. We also showed that the same mechanism of interaction to osteoblasts, adipocytes and chondrocytes that are directly differentiated from the same MSCs. Finally, we have found that the presence of 10% FBS in the infection medium is essential as it helps in achieving the best specific bacterial-cell association with the least background association. The results reveals a mechanism of interaction between *S. aureus* and MSCs that could pave the way for therapeutic intervention that minimises the burden of infection in inflammatory diseases.

## Introduction

Mesenchymal stem cells (MSCs) are a heterogeneous population of cells commonly derived from bone marrow, adipose tissue and placenta (1). The major characteristics of MSCs, are their fibroblast like morphology, their ability to adhere to plastic and their ability to differentiate along the osteogenic, chondrogenic and adipogenic pathways. The mechanism underlying the differentiation process is complex and involves the coordinated action of several cytokines, growth factors and other proteins (2). In addition to their ability to differentiate, MSCs have the ability to repair damaged tissues and possess immunomodulatory properties (3). Human bone marrow derived MSCs migrate to a site of injury, release proinflammatory cytokines and modulate T-cell proliferation following ligand binding to toll like receptor (TLR)3 and 4 (4). At sites of injury, MSCs may well encounter bacterial pathogens. Josse et al, 2014, for example, identified a direct interaction between MSC derived from Wharton’s jelly and the Gram positive pathogen *Staphylococcus aureus* (5). However, the precise molecular details of this interaction remain to be elucidated.

*S. aureus*, is a common cause of focal as well as disseminated diseases. *S. aureus* can enter the blood and from there cause infections of the cardiac valves, endocarditis and bone tissue, causing septic arthritis and osteomyelitis (6). An inflammatory condition of the bone, osteomyelitis can involve the cortical or trabecular bone as well as the periosteum and bone marrow (Add Maffulli et al 2016). Epidemiological data suggest that *S. aureus* is responsible for ~80% of osteomyelitis cases (7). Thus it is plausible the *S aureus* and MSC’s could come into direct contact with each other during osteomyelitis involving the bone marrow.

*S. aureus* possess numerous virulence factors which play a role in the development and progression of disease, through promoting the association and invasion of bacteria, protection of bacteria from host defence mechanisms, and causing damage to tissue during infection (8). A major group of virulence factors produced by *S. aureus* are the Microbial Surface Component Recognising Adhesive Matrix Molecules (MSCRAMM’s). Such molecules play an important role in mediating bacterial interactions with a number of extracellular matrix proteins including, fibronectin, fibrinogen, bone sialoprotein, collagen, and elastin (9–11). Moreover, such interactions allow bacteria to use the extracellular matrix proteins to form a molecular bridge between host cell receptors and bacterial surface proteins (12). *S. aureus* can use MSCRAAMs such as the fibronectin binding proteins (FnBPA and FnBPB), infect a number of mammalian cell type, including epithelial cells (16, 19), endothelial cells (16, 20), fibroblasts (16), and osteoblasts (17, 21–23). Surface proteins such as MSCRAMMs are often implicated in this process and In support of this, a number of studies have demonstrated that the invasion of host cells is poor by *S. aureus* mutants lacking the expression of FnBPs, both or each one alone (16, 17). Arciola *et al*. stated that from a variety of orthopaedic infections, 98% and 99% of clinical isolates possessed *fnbA* and *fnbB* respectively, while only 46% of the isolates express *cna* (a gene encoding collagen binding protein) (18).

In the present study we aim to show if a direct interaction occurs between *S. aureus* and bone marrow derived MSCs and identify if MSCRAMMs have a role in this interaction. We further aim to compare *S. aureus* interaction with cells that differentiate from MSCs, namely; osteoblasts, adipocytes and chondrocytes.

## Materials and Methods

### Isolation and expansion of Mesenchymal stem cells

Bone marrow aspirates were collected into heparinised tubes to avoid samples clotting. Subsequently, 1 ml of each aspirate was dispersed in T175 culture flasks containing 25 ml of low glucose Dulbecco’s Modified Eagle’s Medium (DMEM) (Sigma, Dorset, UK), supplemented with 10% (v/v) FBS, 1% penicillin/streptomycin, 1% (v/v) glutamax, and 10 ng/ml FGF-2. After XXX days, population purity and phenotype were assessed by flow cytometry. Human MSCs at passage 4 were seeded into 96 well plates at a density of 5000 cells/cm^2^ until confluence and used for differentiation or infection experiments as described below.

### Mesenchymal stem cell differentiation

MSCs differentiation was effected using aHuman Mesenchymal Stem Cell Functional Identification Kit, (R&D Systems, Inc.) according to the manufacturer’s instructions. Following differentiation the resulting osteoblasts, adipocytes and chondrocytes were seeded into 96 well plates and grown to confluency for subsequent infection experiments.

### Bacterial Strains

*Staphylococcus aureus* 8325.4 and corresponding *Staphylococcus epidermidis* (used as a negative control for *S. aureus* interaction), as listed in table 1, were obtained from Dr Andrew Edwards, University of Bath.

**Table 1.**
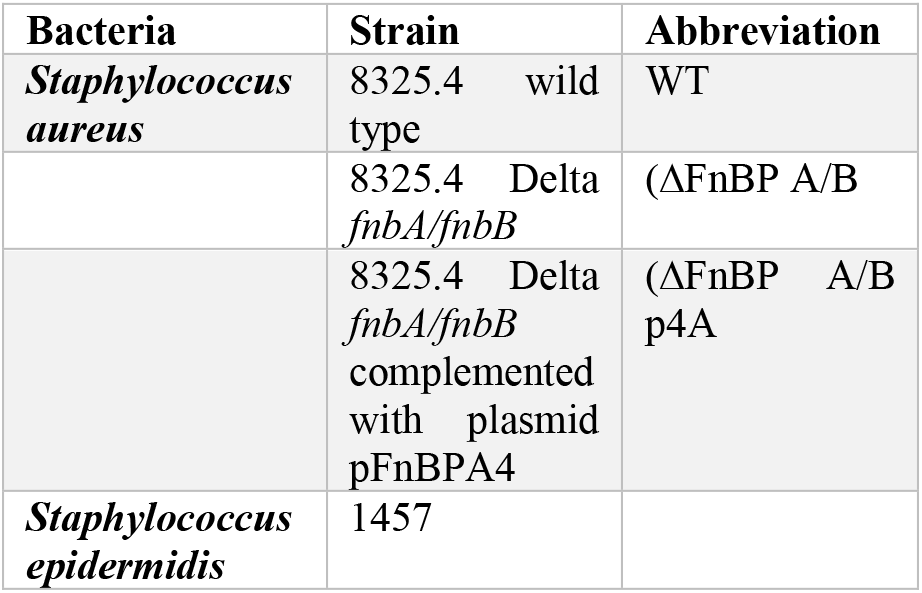
Different bacterial strains used in the experiments.

| Bacteria | Strain | Abbreviation |
| --- | --- | --- |
| *Staphylococcus aureus* | 8325.4 wild type | WT |
|  | 8325.4 Delta *fnbA/fnbB* | (ΔFnBP A/B |
|  | 8325.4 Delta *fnbA/fnbB* complemented with plasmid pFnBPA4 | (ΔFnBP A/B p4A |
| *Staphylococcus epidermidis* | 1457 |  |

All bacteria were cultured on Mueller-Hinton agar (MHA) plates and grown at 37 °C for 16 hours in a humidified incubator with 5% CO_2_.

#### Cell infection assays

*S. aureus* (WT and mutants) and *S. epidermidis* were harvested form MHA agar, resuspended in phosphate buffered saline (PBS) and optical density measured at 600nm. Bacterial suspensions were adjusted to an appropriate multiplicity of infection (MOI) in PBS based upon an optical density of 1 at 600 nm being equivalent to 2×10^9^ colony forming units (CFU)/ml for *S. aureus* and 0.9×10^9^ CFU/ml for *S. epidermidis*.

MSCs in 96 well plate were washed to remove antibiotics and incubated with each bacterial species at a multiplicity of infection (MOI) of 100 in 100 μl of infection medium (DMEM supplemented with glutamax, and FGF-2). In each case cells were infected for 90 min at 37°C in 5% CO_2_. In order to determine the effect of FBS, Human serum and plasma fibronectin on bacterial adhesion, infection medium was supplemented with either 10% FBS, 10% heat inactivated human serum (HuSer), or 400 μg/ml plasma fibronectin (Fn). (comparable to the Fn concentration in human plasma which is 300-400 μg/ml (26)) and low glucose DMEM with 10% decomplemented human serum (HuSer). At the end of each infection assay unbound bacterial cells were removed by washing each well 4 times with appropriate infection media.

#### Immunofluorecscence Microcopy

Cells were then fixed in 2% PFA (paraformaldehyde) overnight at 4°C. Wells were subsequently washed with dH_2_0 for 5 minutes and non-specific binding sites blocked for 1h at room temperature (RT),with 3% bovine serum albumin (BSA)in PBS.. Bacteria were detected using Rabbit anti- *S. aureus* polyclonal antibody (Abd Serotec; 2 μg/ ml in 1% BSA) or Mouse anti- *S. epidermidis* (Thermo scientific; 3 μg/ ml in 1% BSA). MSC’s were labelled with anti-vimentin monoclonal antibody (Sigma; 1 μg/ ml in 1% BSA). Primary antibodies were detected using 1 μg/ ml of anti-rabbit Alexa Fluor 594 or anti-mouse Alexa Fluor 488 in 1%BSA as appropriate (Invitrogen). All antibodies were incubated at room temperature for 1 hr, then wells were washed 3 times for 5 minutes (PBS with 0.05% v/v Tween 20). Finally, 100 μl of 4’,6-Diamidine-2-Phenylindole (DAPI; 1 μg/ ml in PBS) was added to each wells for 30 min to stain nuclei. Plates were stored in PBS containing 0.05% sodium azide. Images were captured using a Hamamatsu digital camera attached to an Olympus IX70 microscope via Image HCI software.

Bacterial quantificationIn order to determine nuber of bacteria associated with each cell type, following washing to remove unbound bacteria, well were incubated with with 1% Saponin for 30min at 37°C. Dilutions were cultured on MHA plates and viable colony forming units (CFU) counted in order to determine the number of bacteria per well. In order to differentiate invasive from associated bacteria prior to addition of saponin, well were incubated with gentamicin (250 μg/ml for 90min at 37°C)in order to kill extracellular bacteria control wells included the addition of cytochalasin D (Conc to add) which prevents uptake of *S. aureus*. Following the incubation with gentamicin, wells were washed 3 times and CFU determined as above.

### Statistical Analysis

All experiments were performed in triplicate using cells from 3 different patients. Data were evaluated using GraphPad Prism software version 6 (GraphPad Software, La Jolla, CA). Data are expressed as means ± SD, unless otherwise specified and one-way ANOVA test was used to determine the statistical significance, followed by Tukey’s Multiple Comparison Test to compare individual test groups. P values of ≤ 0.05 were considered significant.

## Results

### Immunofluorescence adhesion assay

#### Adhesion of *S. aureus* to MSCs and the role of FnBP

*S. aureus* WT was able to adhere to MSCs as shown in Fig. 1(A) and this adhesion was greatly reduced in the absence of FnBP with the use of ΔFnBPA/B mutant bacteria lacking the expression of FnBP as shown in Fig. 1(B), while the bacterial adhesion was greatly enhanced (more than that with WT) with use of *S. aureus* pFnBPA4 mutant (ΔFnBPA/B supplemented with plasmid expressing only the intact FnBP A) as shown in Fig.1 (C). While Fig.1 (D) showd that *S. epidermidis* 1457 fail to adhere to MSCs. The results shown in the figure 1 confirmed adherence of *S. aureus* to MSCs and that the expression of FnBP by bacteria might facilitate this adherence. Also, FnBP A is capable of supporting the adhesion as adherence is improved with the use of *S. aureus* pFnBPA4 mutant.

**Figure 1.**
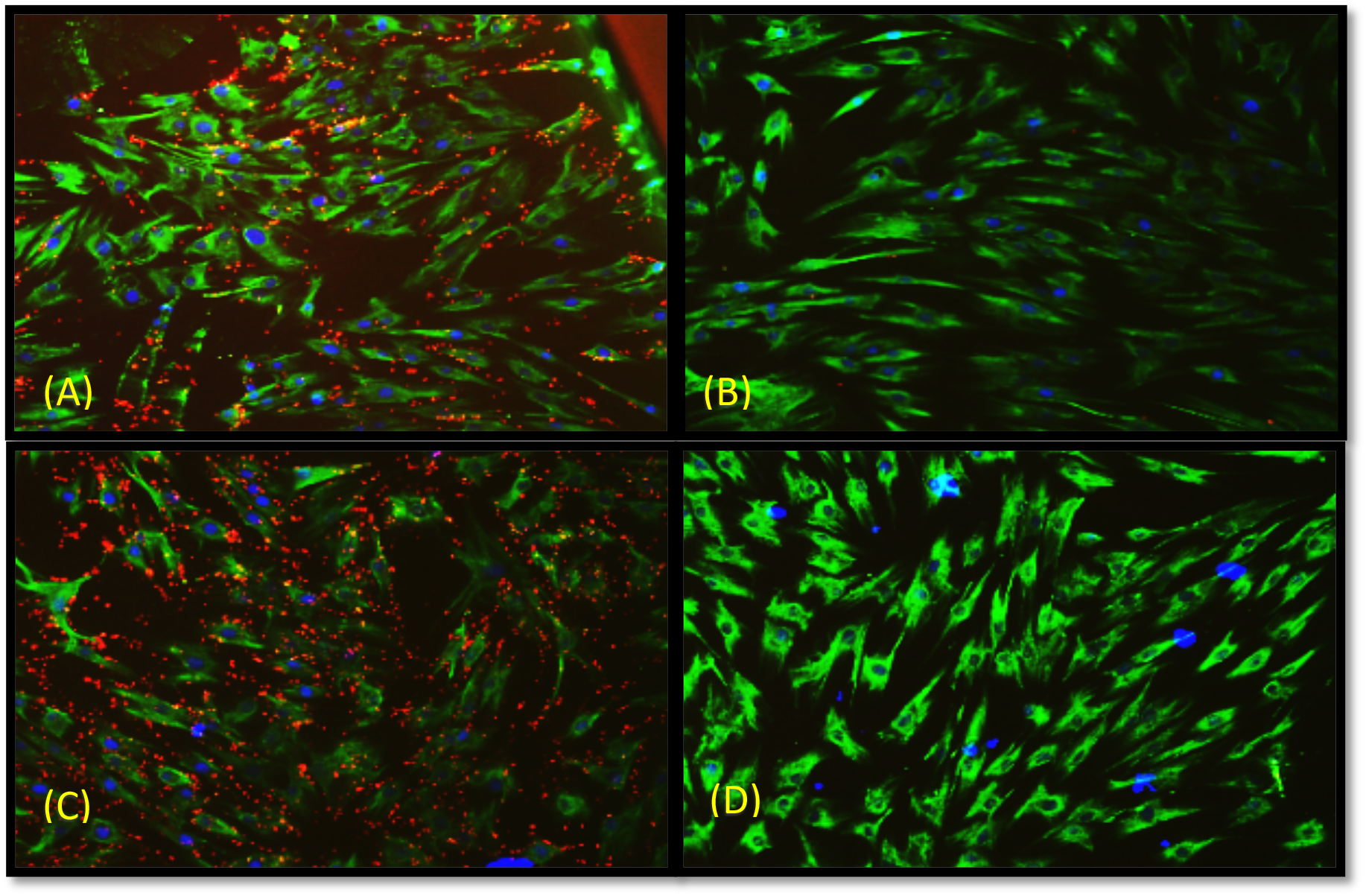
Immunofluorescence adhesion of four bacterial strains to MSCs. The cells were cultured in 96 well plates and then confluent cells were infected with bacterial strains for 90 min. The experiments were repeated with cells from 3 patients and done in technical duplicates. (A) MSCs with S. aureus WT strain. (B) MSCs with S. aureus ΔFnBP A/B Strain. (C) MSCs with S. aureus pFnBPA4 Strain. (D) MSCs with S. epidermidis. Blue areas represent MSCs nucleus stained with DAPI, red dots represent S. aureus bound to rabbit anti S. aureus. Green areas indicate MSCs bound to mouse anti-vimentin. The primary antibodies were detected with secondary antibodies conjugated to the fluorophore.

#### Adhesion of *S. aureus* to differentiated osteoblaB)sts and the role of FnBNPuc

As was the case for MSCs, *S. aureus* WT was capable of adherence to osteoblasts as shown in Fig. 2(A) and that the use of ΔFnBPA/B mutant bacteria lacking the expression of FnBP resulted in great reduction in bacterial adhesion as shown in Fig. 2(B). In comparison, wells with *S. aureus* pFnBPA4 mutant (ΔFnBPA/B supplemented with plasmid expressing only the intact FnBP A (Fig.2 (C)) showed great enhancement in bacterial adhesion (more than that with WT). While Fig.2 (D) showed that very few of *S. epidermidis* 1457 bacteria were able to adhere to osteoblasts in contrast to MSCs were there was no adherence. These results confirmed the ability of *S. aureus* to adhere to osteoblasts which depends largely on the expression of FnBP by the bacteria and that FnBP type supports this adhesion.

**Figure 2.**
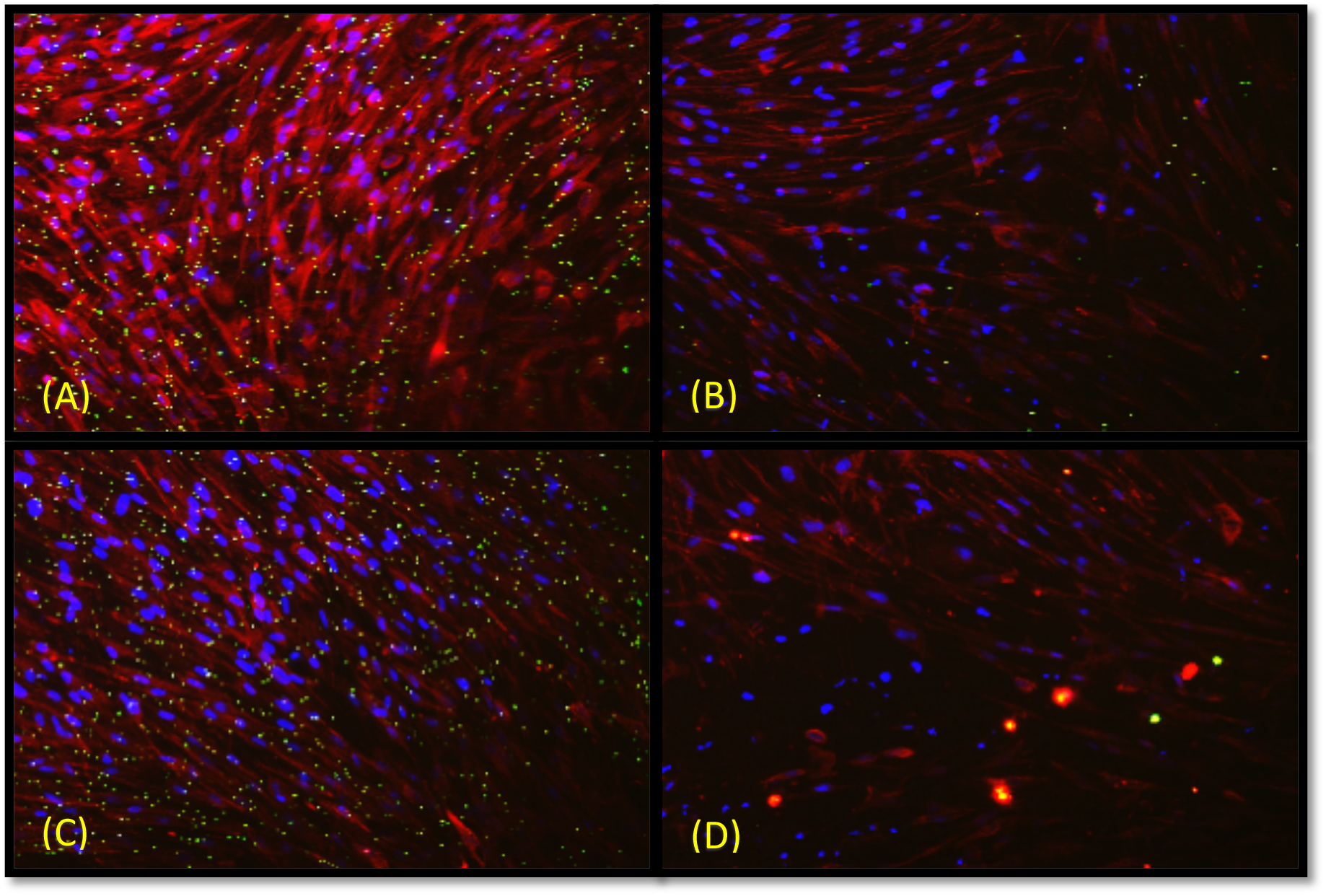
Immunofluorescence adhesion of four bacterial strains to differentiated osteoblasts. After three weeks of MSCs differentiation to osteoblasts in 96 well plate, the differentiated cells were infected with bacterial strains for 90 min. The experiments were repeated with cells from 3 patients and done in technical duplicates. (A) Osteoblasts with S. aureus WT strain. (B) Osteoblasts with S. aureus ΔFnBP A/B Strain. (C) Osteoblasts with S. aureus pFnBPA4 Strain. (D) Osteoblasts with S. epidermidis. Blue areas represent osteoblasts nucleus stained with DAPI, green dots represent S. aureus bound to rabbit anti S. aureus Ab. Red areas indicate osteoblasts bound to mouse anti-osteocalcin Ab. The primary antibodies were detected with secondary antibodies conjugated to the fluorophore.

#### Adhesion of *S. aureus* to differentiated adipocytes and the role of FnBP

The results were comparable to that of osteoblasts regarding the adhesion capabilities of *S. aureus* WT to adipocytes and the great reduction in adhesion with the use of ΔFnBPA/B mutant bacteria which is restored again and to a higher degree with the use of *S. aureus* pFnBPA4 mutant as shown in Fig.3 (A), (B), and (C). Also, Fig.3 (D) showed similar trend for *S. epidermidis* 1457 with very few bacterial adherence. This states the importance of *S. aureus* FnBP expression in their adherence to adipocytes.

**Figure 3.**
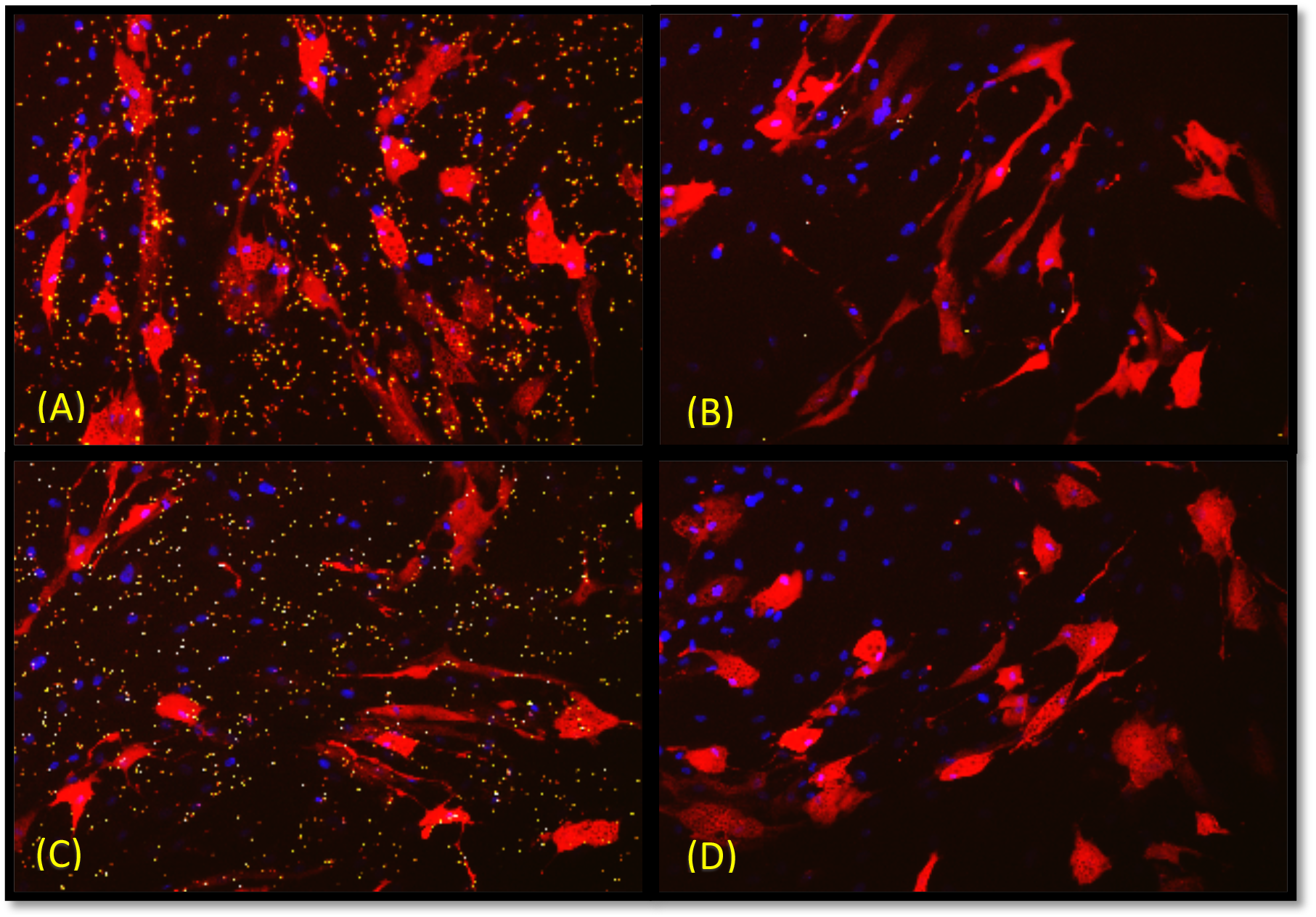
Immunofluorescence adhesion of four bacterial strains to differentiated adipocytes. After three weeks of MSCs differentiation to adipocytes in 96 well plate, the differentiated cells were infected with bacterial strains for 90 min. The experiments were repeated with cells from 3 patients and done in technical duplicates. (A) Adipocytes with S. aureus WT strain. (B) Adipocytes with S. aureus ΔFnBP A/B strain. (C) Adipocytes with S. aureus pFnBPA4 Strain. (D) Adipocytes with S. epidermidis. Blue areas represent adipocytes nucleus stained with DAPI, green-yellow dots represent S. aureus bound to rabbit anti S. aureus Ab. Red areas indicate adipocytes bound to goat anti-fatty acid binding protein Ab. The primary antibodies were detected with secondary antibodies conjugated to the fluorophore.

#### Adhesion of *S. aureus* to differentiated chondrocytes and the role of FnBP

The results shown in Fig.4 (A), (B), and (C) confirmed that *S. aureus* WT was able to adhere to chondrocytes and the absence of FnBP (with the use of ΔFnBPA/B mutant bacteria) greatly affected the adhesion in a negative way, while the use of *S. aureus* pFnBPA4 mutant greatly enhanced adhesion capability of *S. aureus*, even to a higher degree that WT strain. Also, *S. epidermidis* 1457 strain was unable to adhere to chondrocytes as shown in Fig.4 (D). These results stated a comparable findings to that of *S. aureus* interaction with osteoblasts and adipocytes in regard to the importance of FnBP in bacterial-cell interaction and the enhancement of bacterial adhesion.

**Figure 4.**
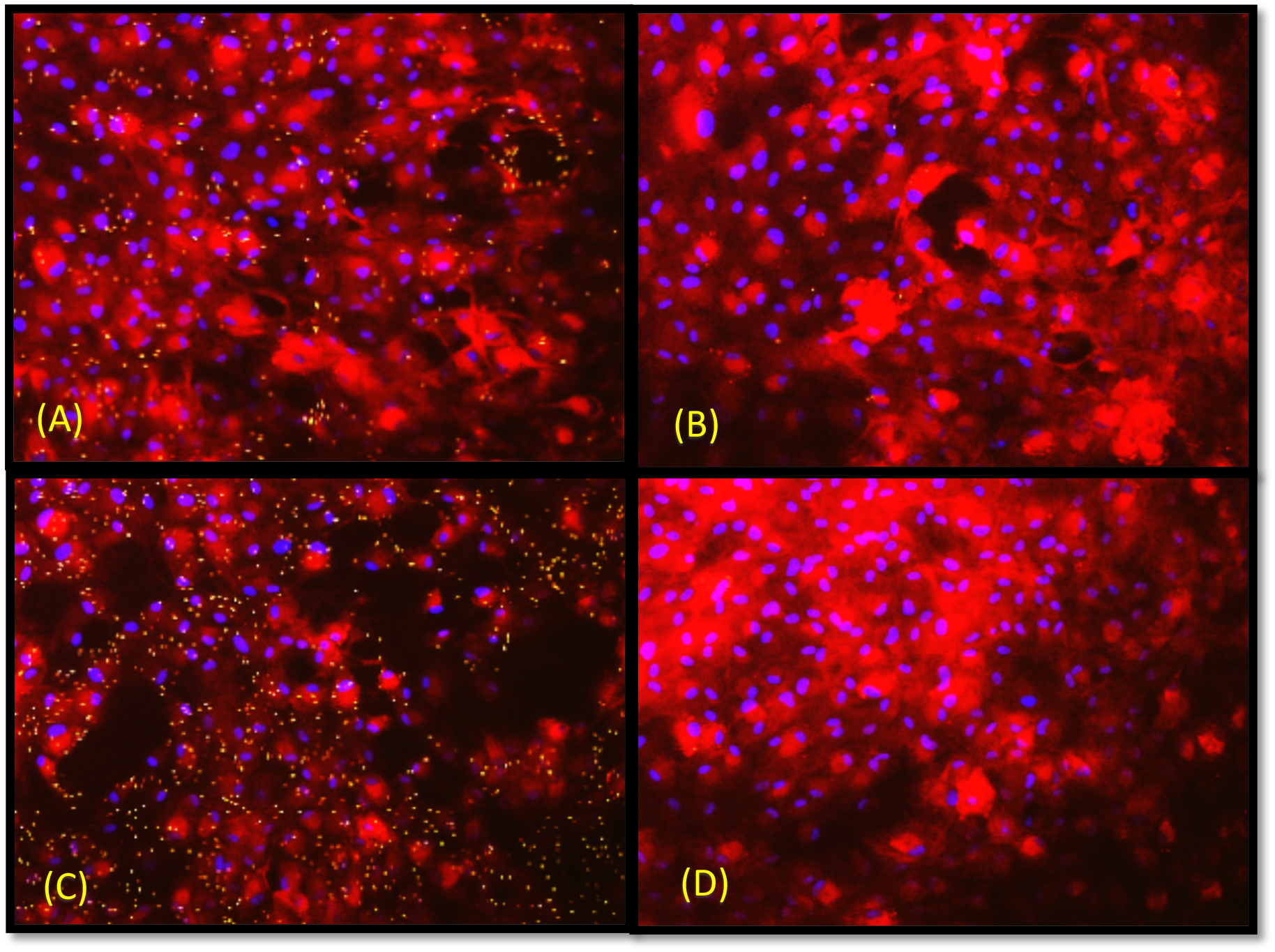
Immunofluorescence adhesion of four bacterial strains to differentiated chondrocytes. After three weeks of MSCs differentiation to chondrocytes in 96 well plate, the differentiated cells were infected with bacterial strains for 90 min. The experiments were repeated with cells from 3 patients and done in technical duplicates. (A) Chondrocytes with *S. aureus* WT strain. (B) Chondrocytes with *S. aureus* ΔFnBP A/B Strain. (C) Chondrocytes with *S. aureus* pFnBPA4 Strain. (D) Chondrocytes with *S. epidermidis*. Blue areas represent chondrocytes nucleus stained with DAPI, green dots represent *S. aureus* bound to rabbit Anti *S. aureus* Ab. Red areas indicate chondrocytes bound to goat anti-aggrecan Ab. The primary antibodies were detected with secondary antibodies conjugated to the fluorophore.

#### The influence of FBS, Human Serum and plasma fibronectin on bacterial adhesion

The highest bacterial adhesion was seen between MSCs and *S. aureus* WT in the presence of 10% FBS with least bacterial association control as shown Fig. 5 (E) and (F) and this adhesion greatly reduced with the use of mutant strain ΔFnBP in the same conditions, Fig. 5 (G) and (H). The wells with low glucose DMEM without serum show large bacterial number of both strains, WT and ΔFnBP, but with nonspecific cell adhesion (mostly in the background of the well), Fig. 5 (A) and (C) as confirmed with the bacterial association wells, Fig. 5 (B) and (D). The use of 40 mg/ml of plasma fibronectin result in great reduction in bacterial adhesion with the MSCs for WT strain with non-specific adhesion for mutant strain (mostly background adhesion), Fig. 5 (I) and (K), with high bacterial association control, comparable to the wells without serum, Fig. 5 (J) and (L). Lastly, the addition of 10% decomplemented human serum seems to greatly reduce bacterial number in the wells in the presence and absence of cells for both bacterial strains, Fig. 5 (M),(N),(O),(P), with some bacterial clumps for WT strain as shown in Fig. (M).

**Figure 5.**
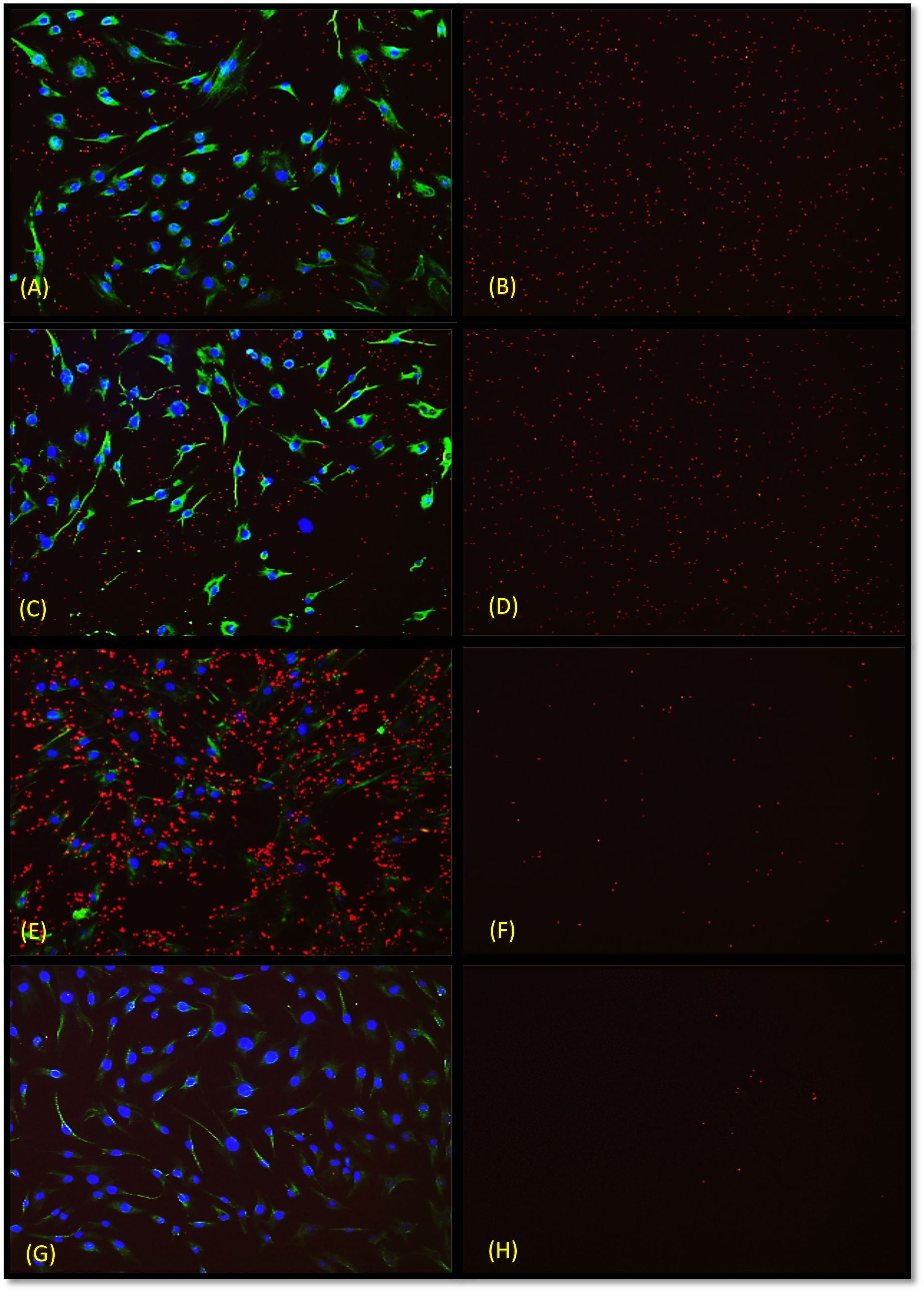
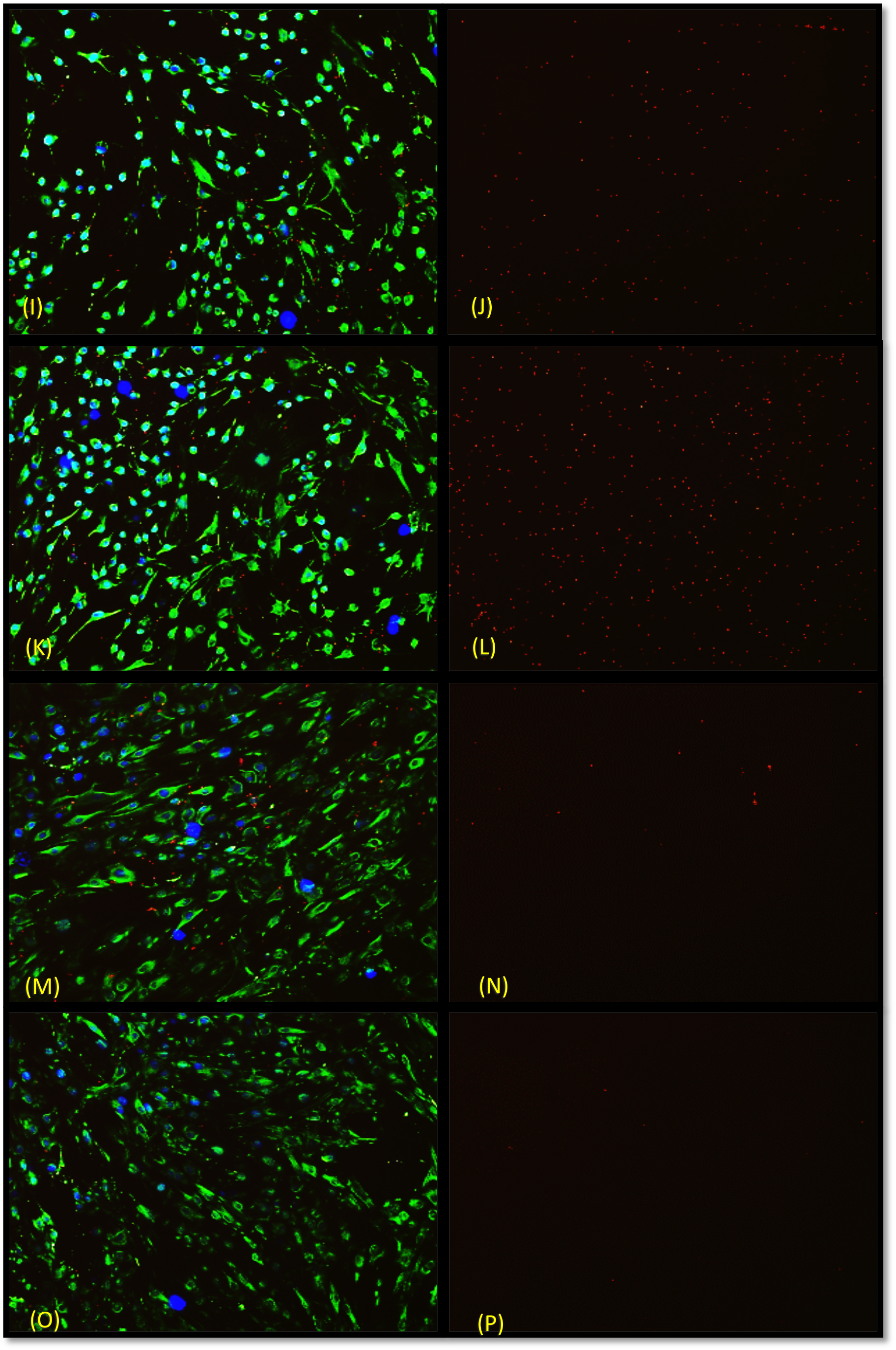
Effect of FBS, Fn, and HuSer on bacterial adhesion to MSCs. Infection assay was done in 96 well plate using 2 bacterial strains with MSCs in 4 different conditions for 90 min, then washing and immunohistochemical staining was done, these are representative images from triplet for three patients. (A) MSCs with *S. aureus* WT in low glucose DMEM without serum (B) *S. aureus* WT association control in low glucose DMEM without serum (C) MSCs with *S. aureus* ΔFnBP in low glucose DMEM without serum (D) *S. aureus* ΔFnBP association control in low glucose DMEM without serum (E) MSCs with *S. aureus* WT in low glucose DMEM + 10% FBS (F) *S. aureus* WT association control in low glucose DMEM + 10% FBS (G) MSCs with *S. aureus* ΔFnBP in low glucose DMEM +10% FBS (H) *S. aureus* ΔFnBP association control in low glucose DMEM +10%FBS (I) MSCs with *S. aureus* WT in low glucose DMEM + 400 ìg/ml Fn (J) *S. aureus* WT association control in low glucose DMEM + 400 ìg/ml Fn (K) MSCs with *S. aureus* ΔFnBP in low glucose DMEM + 400 ìg/ml Fn (L) *S. aureus* ΔFnBP association control in low glucose DMEM + 400 ìg/ml Fn (M) MSCs with *S. aureus* WT in low glucose DMEM + 10% HuSer (N) *S. aureus* WT association control in low glucose DMEM + 10% HuSer (O) MSCs with *S. aureus* ΔFnBP in low glucose DMEM + 10% HuSer (P) *S. aureus* ΔFnBP association control in low glucose DMEM + 10%HuSer. Blue areas represent MSCs nucleus stained with DAPI, red dots represent *S. aureus* bound to rabbit anti *S. aureus* Ab. Green areas indicate MSCs bound to mouse anti-vimentin Ab. The primary antibodies were detected with secondary antibodies conjugated to the fluorophore.

#### Association and Invasion Assay for MSCs

Fig. 6 demonstrate that 14.66% of *S. aureus* WT CFU that present in the well associated with MSCs which is statistically higher (p value ≤ 0.001) than that for *S. aureus* ΔFnBP (0.23%) and *S. epidermidis* (0.32%), while *S. aureus* ΔFnBP p4A, a mutant strain expressing intact FnBP type A have the highest association at 40.26%, which is statistically higher than WT strain (p value ≤ 0.001). These results confirm the ability of *S. aureus* to associate with MSCs and that FnBP plays a major role in this association, while the increase in association with the use ΔFnBP p4A confirm the role of FnBP in the association and that Type A FnBP has a higher significance.

**Figure 6.**
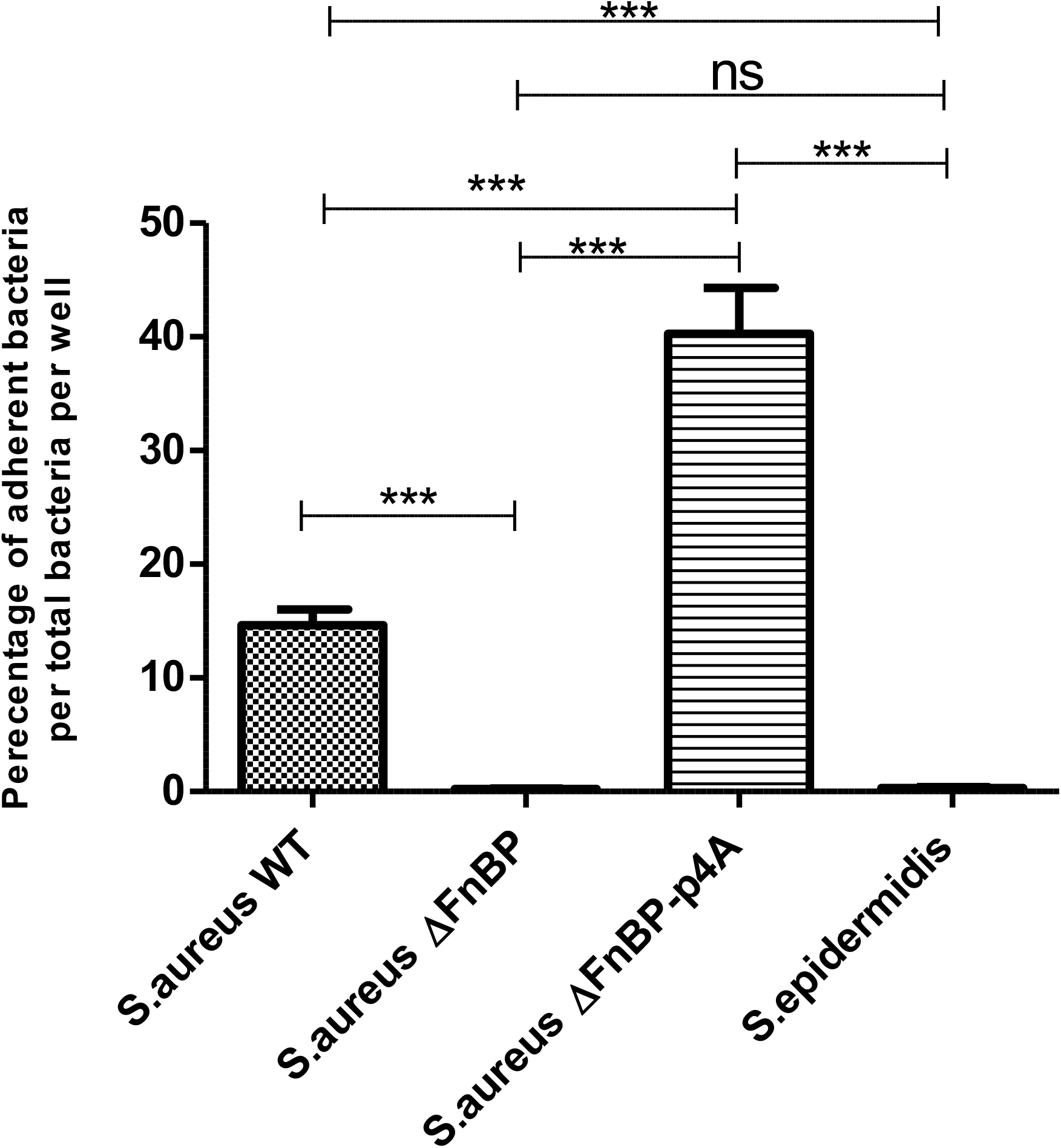
Association assay for MSCs with 4 bacterial strains. Conflunt MSCs in 96 well plate were infected with 4 bacterial strains at MOI 100 for 3 hr, then washing and serial dilution was done followed by inoculation of dilution factors in MHA plates and CFU count done after 24 hr. The data are from three independent experiments of three patients. WT=wild type, DFnBP=mutant lacking the expression of fibronectin binding protein A/B, DFnBP p4A=mutant supplemented with plasmid expressing intact fibronectin binding protein type A. *** = P < 0.001, ns = P > 0.05

After confirming bacterial association with MSCs, extracellular bacteria were killed by gentamicin in order to prove the intracellular presence of *S. aureus* in the MSCs. The results stated that for *S. aureus* WT, 1.59% of the associated bacteria invaded MSCs, which is higher, with a high statistical significance (p value ≤ 0.001), than that for *S. aureus* ΔFnBP (0.47%) and *S. epidermidis* (0.55%) as shown in Fig. 14, while ΔFnBP p4A show higher invasion (2.05%) than other strains with slight statistical difference from *S. aureus* WT (p value ≤ 0.05) as shown in Fig. 7. These results show the ability of *S. aureus* to invade MSCs and that the invasion is greatly reduced in the absence of FnBP, while the use of ΔFnBP p4A, a mutant strain expressing intact FnBP type A, greatly enhance the invasion to a degree slightly higher the wild type strain.

**Figure 7.**
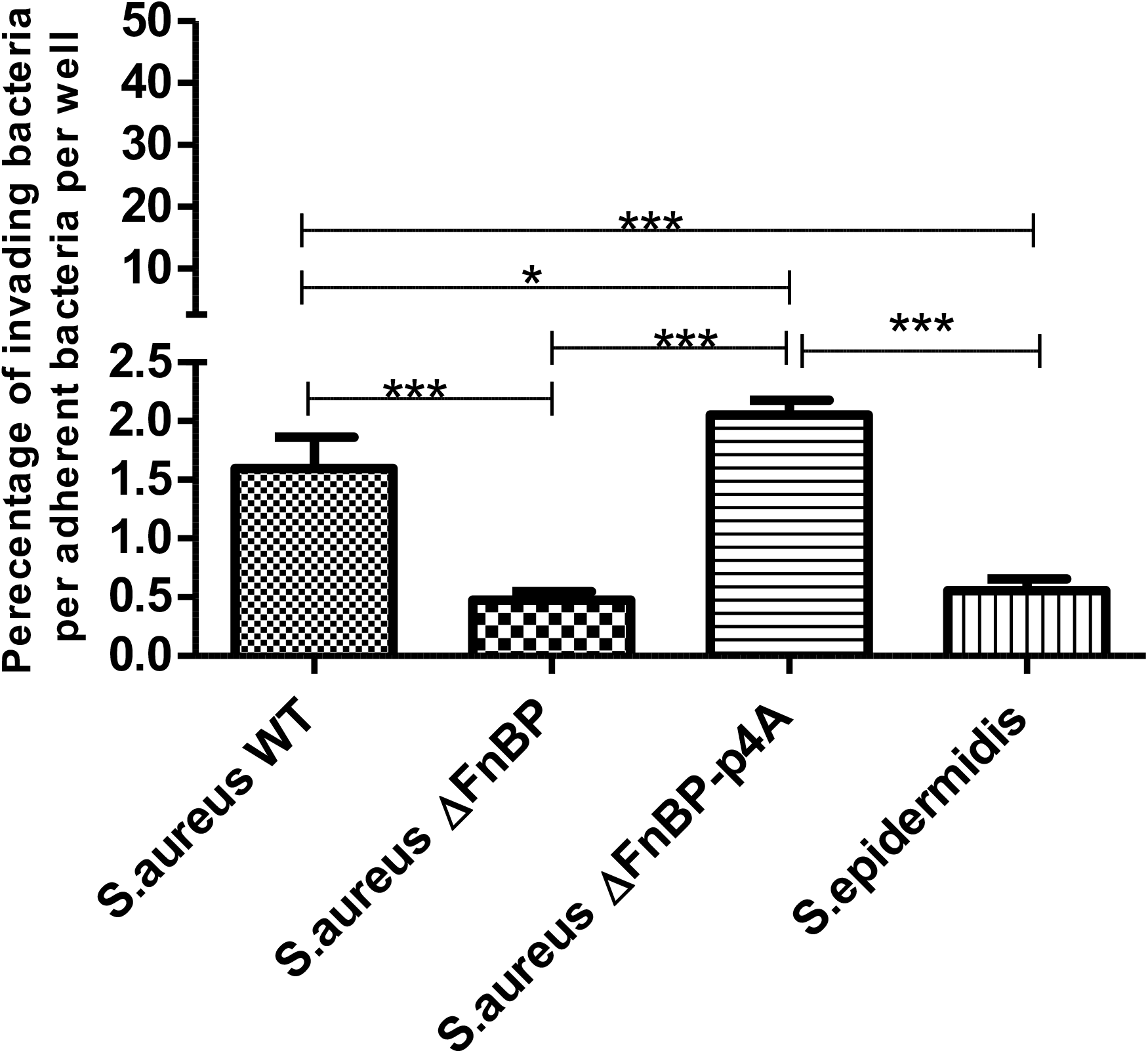
Invasion assay for MSCs with 4 bacterial strains. Conflunt MSCs in 96 well plate were infected with 4 bacterial strains at MOI 100 for 3 hr, then washing and the plate incubated with gentamycin for 90 min to kill extracellular bacteria, then washing and serial dilution was done followed by inoculation of dilution factors in MHA plates and CFU count done after 24 hr. The data are from three independent experiments of three patients. WT=wild type, DFnBP=mutant lacking the expression of fibronectin binding protein A/B, DFnBP p4A=mutant supplemented with plasmid expressing intact fibronectin binding protein type A. *** = P < 0.001, * = P < 0.05

#### Association and Invasion Assay for differentiated Osteoblasts

Fig. 8 demonstrate that 13.75% of *S. aureus* WT CFU that present in the well associated with osteoblasts which is statistically higher (p value ≤ 0.001) than that for *S. aureus* ΔFnBP (1.6%) and *S. epidermidis* (2.82%), while *S. aureus* ΔFnBP p4A, a mutant strain expressing intact FnBP type A have the highest association at 20.76%, which is statistically higher than WT strain (p value ≤ 0.05). These results confirm the ability of *S. aureus* to associate with osteoblasts and that FnBP plays a major role in this association, while the increase in association with the use ΔFnBP p4A confirm the role of FnBP in the association and that Type A FnBP has a higher significance.

**Figure 8.**
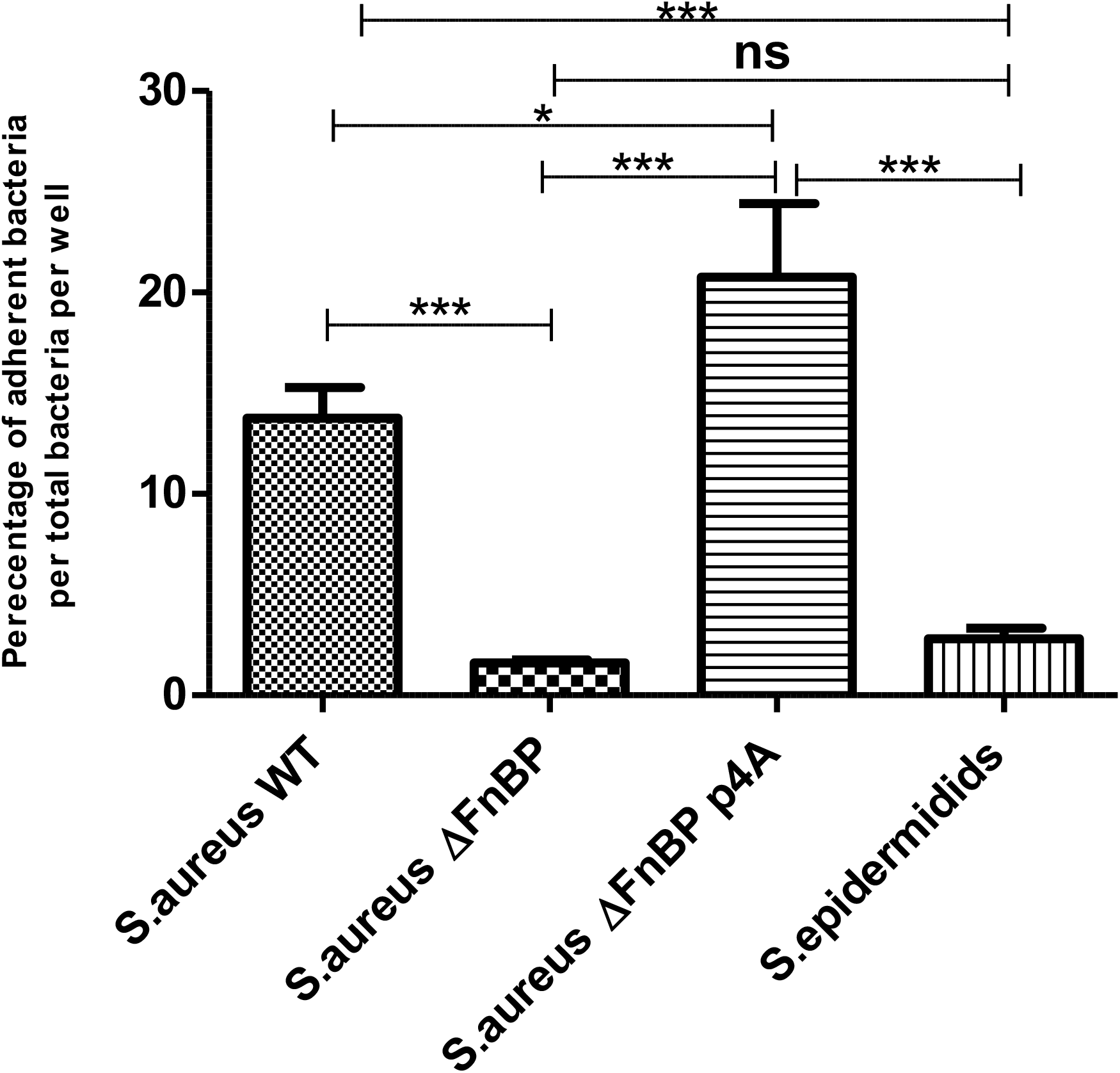
Association assay for osteoblasts with 4 bacterial strains. Conflunt osteoblasts in 96 well plate were infected with 4 bacterial strains at MOI 100 for 3 hr, then washing and serial dilution was done followed by inoculation of dilution factors in MHA plates and CFU count done after 24 hr. The data are from three independent experiments of three patients. WT=wild type, DFnBP=mutant lacking the expression of fibronectin binding protein A/B, DFnBP p4A=mutant supplemented with plasmid expressing intact fibronectin binding protein type A. *** = P < 0.001, * = P < 0.05, ns = P > 0.05

After confirming bacterial association with osteoblasts, extracellular bacteria were killed by gentamicin in order to prove the intracellular presence of *S. aureus* in the osteoblasts. The results stated that for *S. aureus* WT, 1.36% of the associated bacteria invaded osteoblasts, which is higher, with a high statistical significance (p value ≤ 0.001), than that for *S. aureus* ΔFnBP (0.18%) and *S. epidermidis* (0.38%) as shown in Fig. 9, while ΔFnBP p4A show higher invasion (1.60%) than other strains, but with no statistical difference from *S. aureus* WT (p value ≥ 0.5) as shown in Fig. 9. These results show the ability of *S. aureus* to invade osteoblasts and that the invasion is greatly reduced in the absence of FnBP, while the use of ΔFnBP p4A, a mutant strain expressing intact FnBP type A, greatly enhance the invasion to a degree comparable to that of the wild type strain.

**Figure 9.**
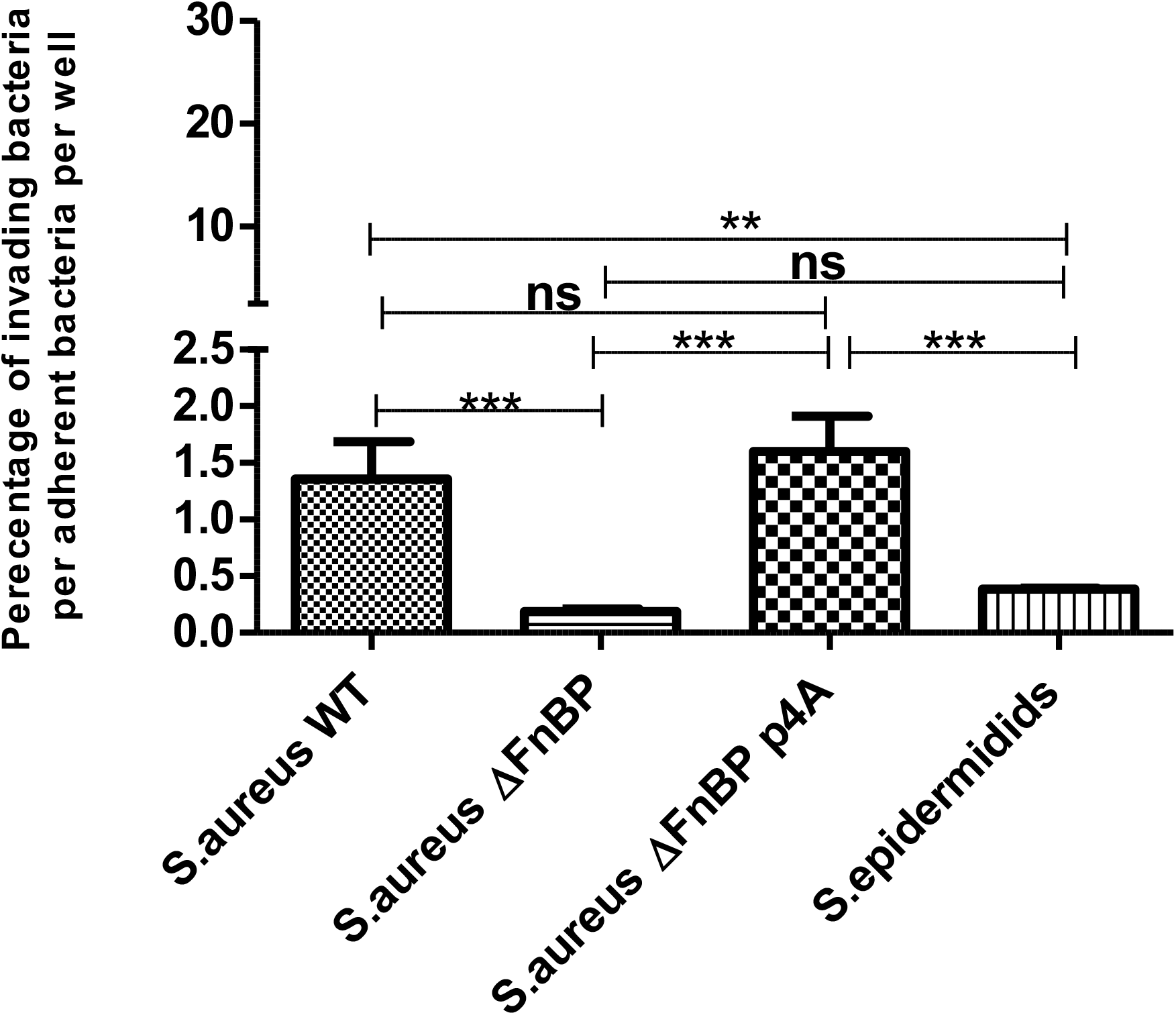
Invasion assay for Osteoblasts with 4 bacterial strains. Conflunt Osteoblasts in 96 well plate were infected with 4 bacterial strains at MOI 100 for 3 hr, then washing and the plate incubated with gentamycin for 90 min to kill extracellular bacteria, then washing and serial dilution was done followed by inoculation of dilution factors in MHA plates and CFU count done after 24 hr. The data are from three independent experiments of three patients. WT=wild type, DFnBP=mutant lacking the expression of fibronectin binding protein A/B, DFnBP p4A=mutant supplemented with plasmid expressing intact fibronectin binding protein type A. *** = P < 0.001, ** = P < 0.01, ns = P > 0.05

#### Association and Invasion Assay for differentiated Adipocytes

Fig. 10 demonstrate that 16.87% of *S. aureus* WT CFU that present in the well associated with Adipocytes which is statistically higher (p value ≤ 0.001) than that for *S. aureus* ΔFnBP (1.44%) and *S. epidermidis* (0.57%), while *S. aureus* ΔFnBP p4A, a mutant strain expressing intact FnBP type A have the highest association at 26.08%, which is statistically higher than WT strain (p value ≤ 0.05). These results confirm the ability of *S. aureus* to associate with Adipocytes and that FnBP plays a major role in this association, while the increase in association with the use ΔFnBP p4A confirm the role of FnBP in the association and that Type A FnBP has a higher significance.

**Figure 10.**
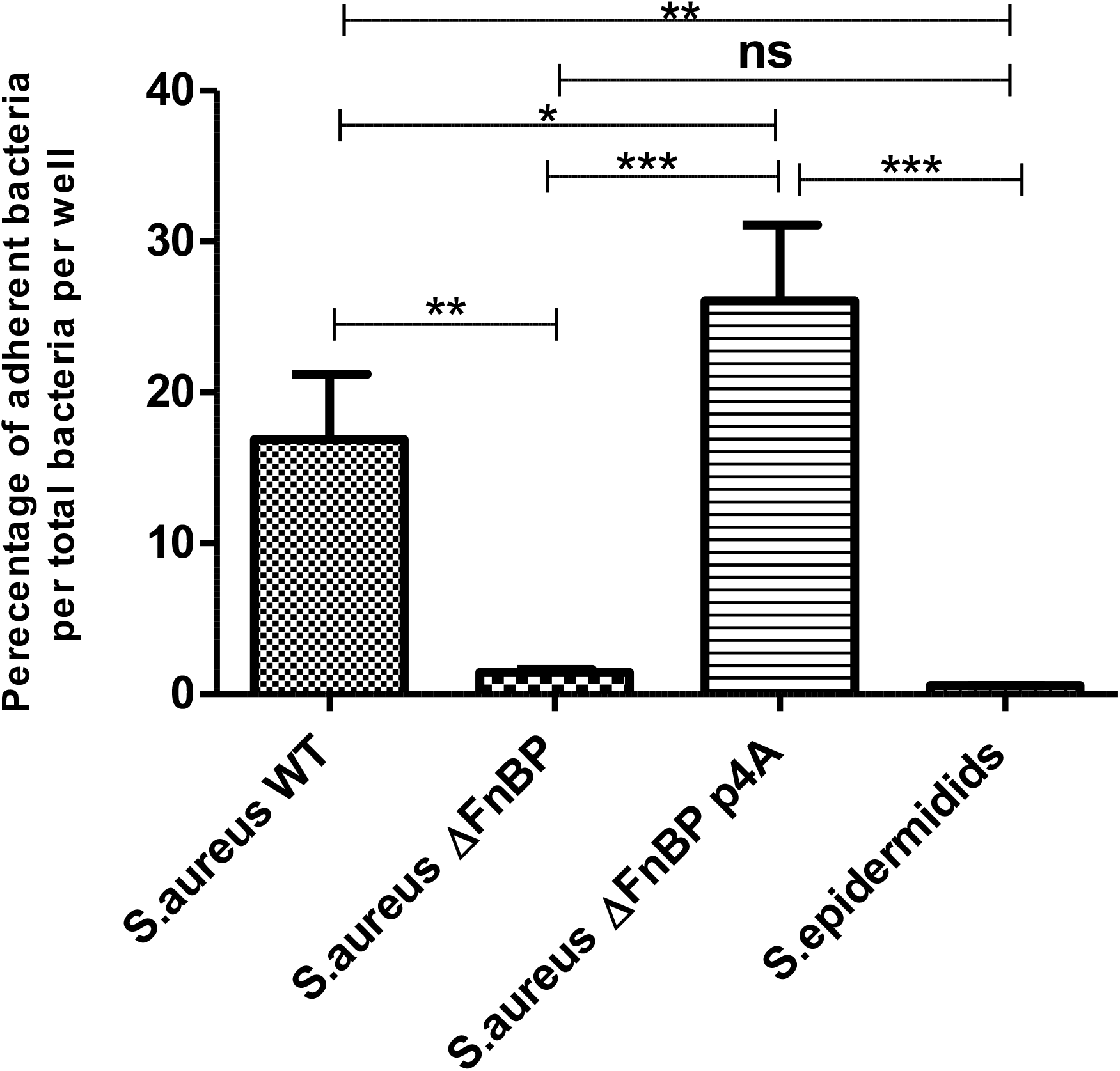
Association assay for adipocytes with 4 bacterial strains. Conflunt adipocytes in 96 well plate were infected with 4 bacterial strains at MOI 100 for 3 hr, then washing and serial dilution was done followed by inoculation of dilution factors in MHA plates and CFU count done after 24 hr. The data are from three independent experiments of three patients. WT=wild type, DFnBP=mutant lacking the expression of fibronectin binding protein A/B, DFnBP p4A=mutant supplemented with plasmid expressing intact fibronectin binding protein type A. *** = P < 0.001, ** = P < 0.01, *= P £ 0.05, ns = P > 0.05

After confirming bacterial association with Adipocytes, extracellular bacteria were killed by gentamicin in order to prove the intracellular presence of *S. aureus* in the Adipocytes. The results stated that for *S. aureus* WT, 1.82% of the associated bacteria invaded Adipocytes, which is higher than that for *S. aureus* ΔFnBP (0.57%) and *S. epidermidis* (0.32%) as shown in Fig. 11 with a statistical significance (p value ≤ 0.001 and ≤ 0.001 respectively). The bacterial invasion increase with the use of bacterial strain ΔFnBP p4A at 1.23%, but there was no statistical significance from the invasion for *S. aureus* WT and ΔFnBP (p value > 0.5) as shown in Fig. 11. These results show the ability of *S. aureus* to invade Adipocytes and that the invasion is greatly reduced in the absence of FnBP, while the use of ΔFnBP p4A, a mutant strain expressing intact FnBP type A, enhance the invasion in slight degree which may indicate that invasion of adipocytes by *S. aureus* may require the expression of both types A and B of the FNBP.

**Figure 11.**
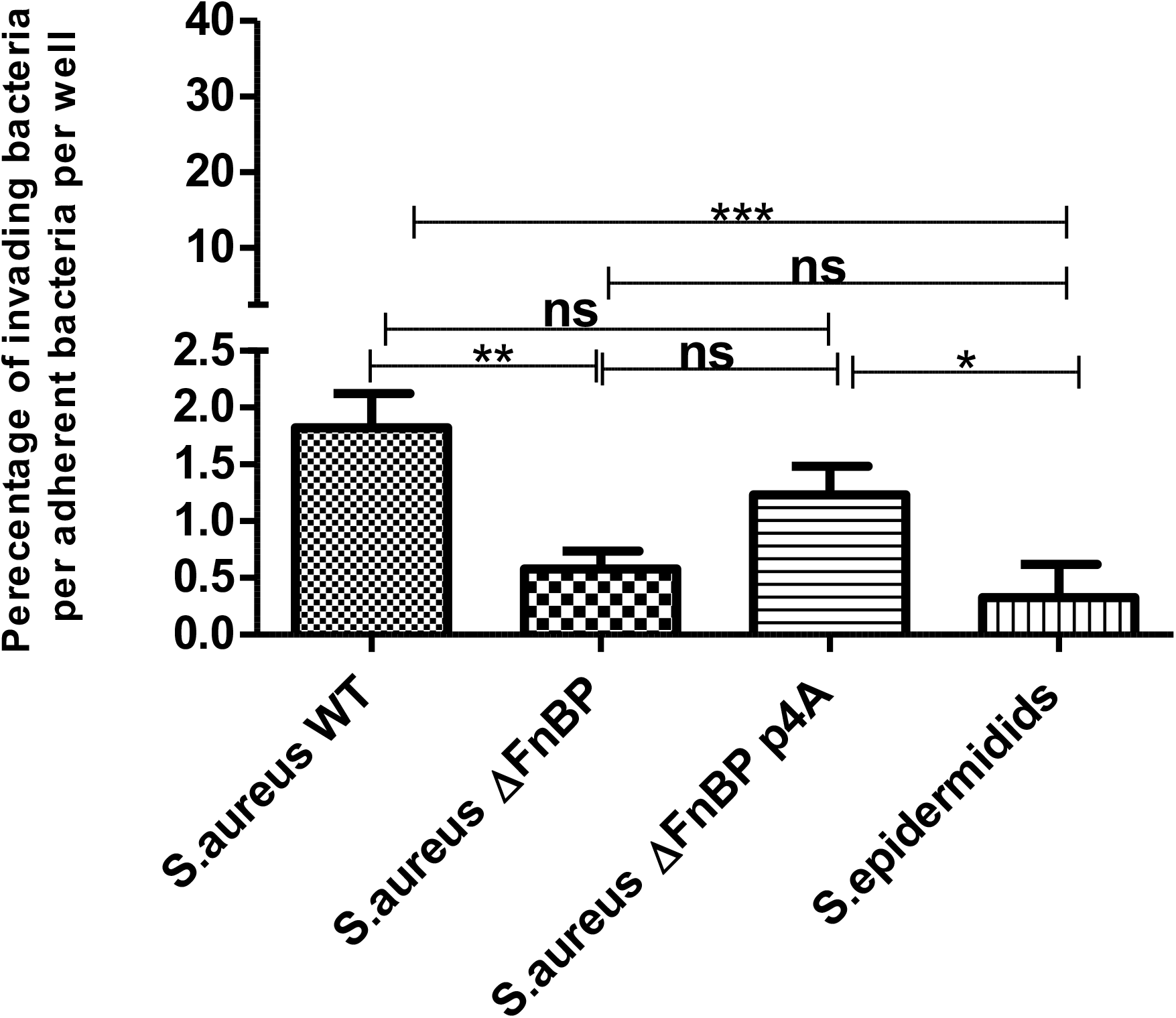
Invasion assay for Adipocytes with 4 bacterial strains. Conflunt Adipocytes in 96 well plate were infected with 4 bacterial strains at MOI 100 for 3 hr, then washing and the plate incubated with gentamycin for 90 min to kill extracellular bacteria, then washing and serial dilution was done followed by inoculation of dilution factors in MHA plates and CFU count done after 24 hr. The data are from three independent experiments of three patients. WT=wild type, DFnBP=mutant lacking the expression of fibronectin binding protein A/B, DFnBP p4A=mutant supplemented with plasmid expressing intact fibronectin binding protein type A. *** = P < 0.001, ** = P < 0.01, *= P < 0.05, ns = P > 0.05

#### Association and Invasion Assay for differentiated Chondrocytes

Fig. 12 demonstrate that 9.36% of *S. aureus* WT CFU that present in the well associated with Chondrocytes which is statistically higher (p value ≤ 0.001) than that for *S. aureus* ΔFnBP (0.83%) and *S. epidermidis* (0.54%), while *S. aureus* ΔFnBP p4A, a mutant strain expressing intact FnBP type A have the highest association at 10.43%, which is statistically higher than WT strain (p value ≤ 0.01). These results confirm the ability of *S. aureus* to associate with Chondrocytes and that FnBP plays a major role in this association, while the increase in association with the use ΔFnBP p4A confirm the role of FnBP in the association and that Type A FnBP has a higher significance.

**Figure 12.**
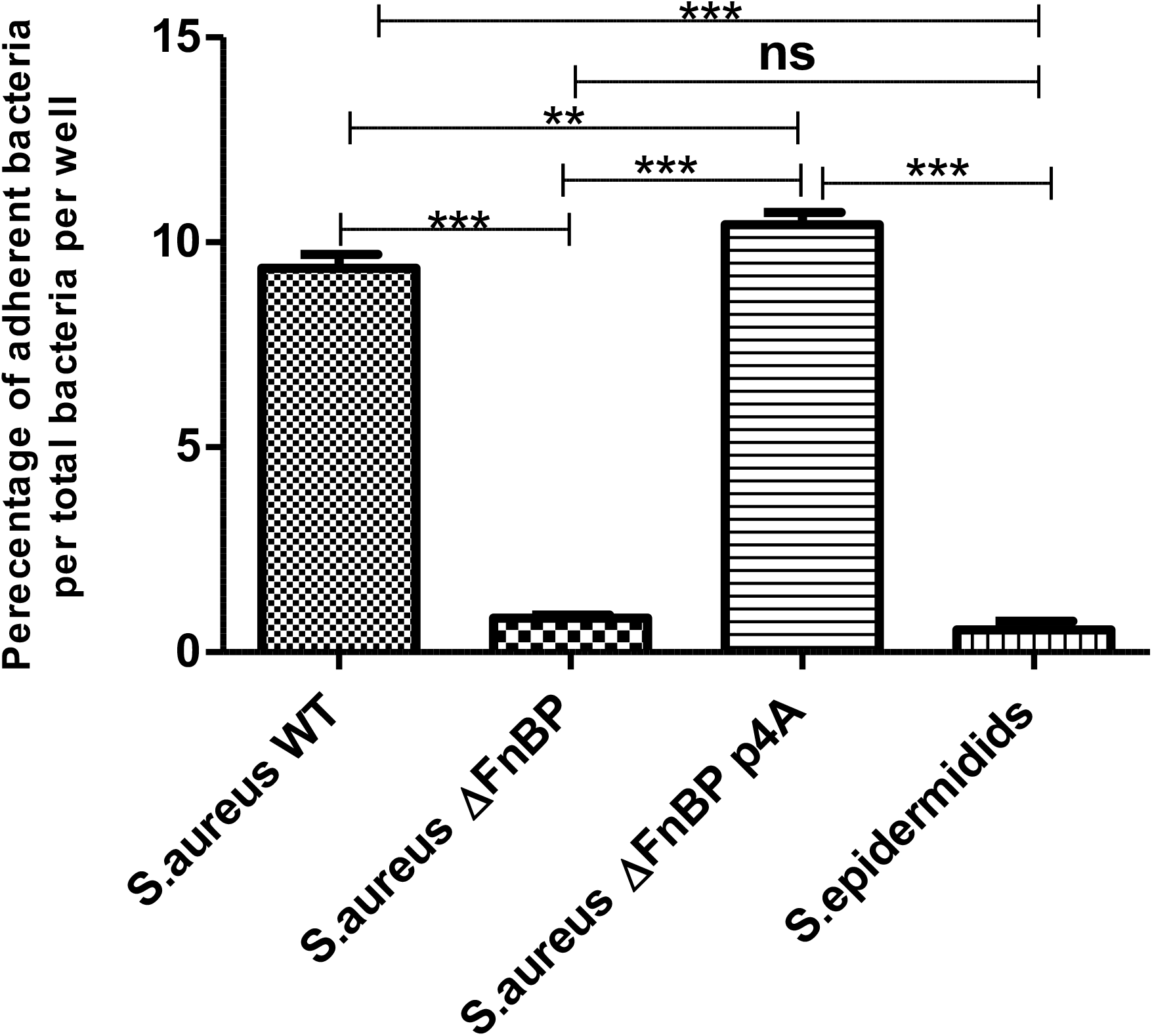
Association assay for chondrocytes with 4 bacterial strains. Conflunt chondrocytes in 96 well plate were infected with 4 bacterial strains at MOI 100 for 3 hr, then washing and serial dilution was done followed by inoculation of dilution factors in MHA plates and CFU count done after 24 hr. The data are from three independent experiments of three patients. WT=wild type, DFnBP=mutant lacking the expression of fibronectin binding protein A/B, DFnBP p4A=mutant supplemented with plasmid expressing intact fibronectin binding protein type A. *** = P < 0.001, ** = P < 0.01, ns = P > 0.05

After confirming bacterial association with Chondrocytes, extracellular bacteria were killed by gentamicin in order to prove the intracellular presence of *S. aureus* in the Chondrocytes. The results stated that for *S. aureus* WT, 2.65% of the associated bacteria invaded Chondrocytes, which is higher, with a high statistical significance (p value ≤ 0.001), than that for *S. aureus* ΔFnBP (0.50%) and *S. epidermidis* (0.61%) as shown in Fig. 13, while ΔFnBP p4A show higher invasion (1.98%) than other strains with no statistical difference from *S. aureus* WT (p value >0.5) as shown in Fig. 13. These results show the ability of *S. aureus* to invade Chondrocytes and that the invasion is greatly reduced in the absence of FnBP, while the use of ΔFnBP p4A, a mutant strain expressing intact FnBP type A, enhance the invasion in slight degree which may indicate that invasion of adipocytes by *S. aureus* may require the expression of both types A and B of FnBP.

**Figure 13.**
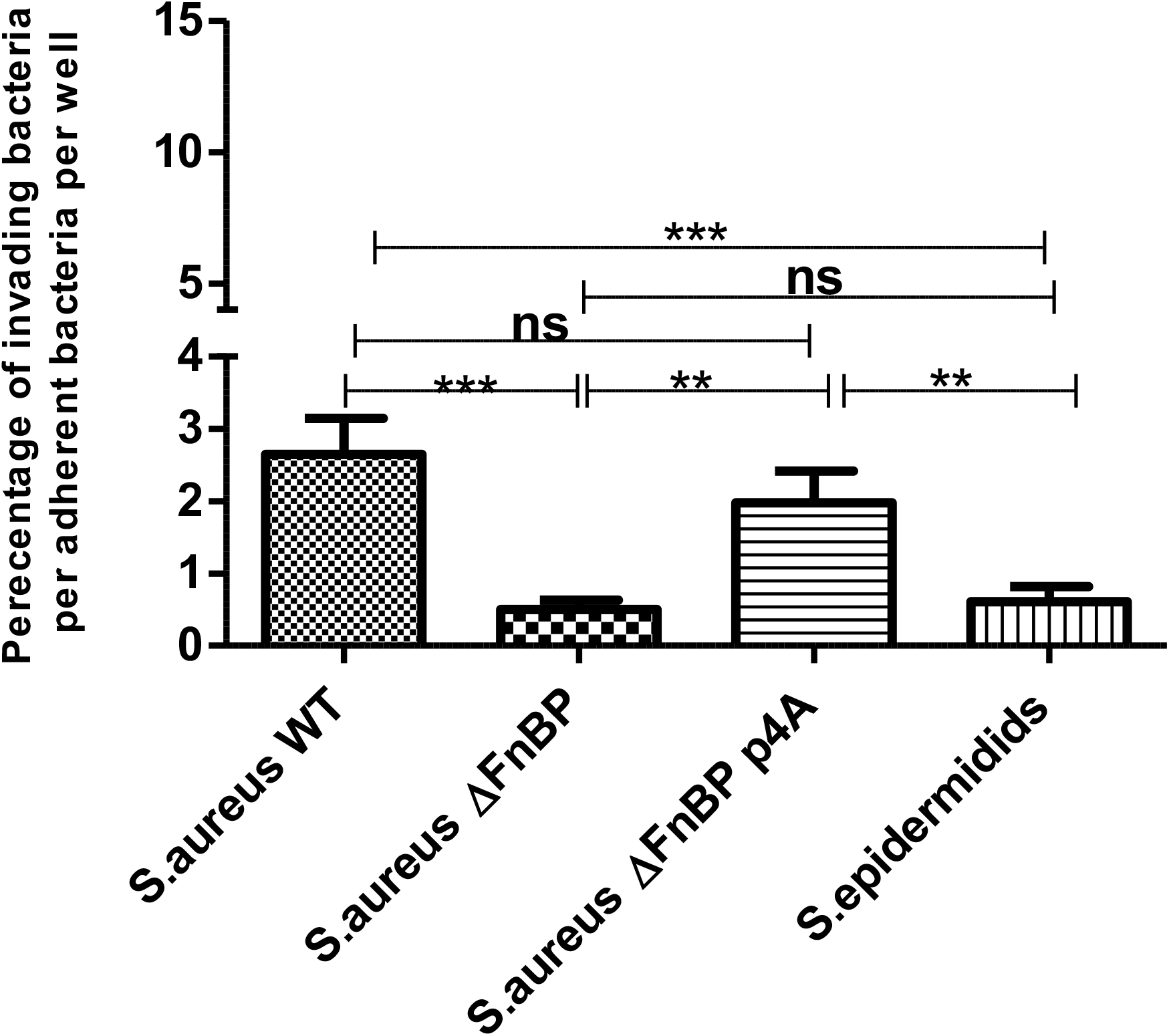
Invasion assay for Chondrocytes with 4 bacterial strains. Conflunt chondrocytes in 96 well plate were infected with 4 bacterial strains at MOI 100 for 3 hr, then washing and the plate incubated with gentamycin for 90 min to kill extracellular bacteria, then washing and serial dilution was done followed by inoculation of dilution factors in MHA plates and CFU count done after 24 hr. The data are from three independent experiments of three patients. WT=wild type, DFnBP=mutant lacking the expression of fibronectin binding protein A/B, DFnBP p4A=mutant supplemented with plasmid expressing intact fibronectin binding protein type A. *** = P < 0.001, ** = P < 0.01, ns = P > 0.05

## DISCUSSION

Until now, very few studies have addressed the interaction between MSCs and *S. aureus* (27–31), but none have studied the interaction of *S. aureus* with bone marrow MSCs and their differentiated osteoblasts, adipocytes and chondrocytes and the possible mechanism of this interaction.

The first aim for the experiments conducted in this study was to confirm the interaction between bone marrow MSCs and *S. aureus*, determining the underlying mechanism and ability of bacteria to invade these cells and survive intracellulary. Then we tried to confirm that this interaction and subsequent invasion and intracellular survival of bacteria, extends to involve the cells which are directly differentiated from these MSCs, namely osteoblasts, adipocytes and chondrocytes.

Infection assay was done using MOI 100 as preliminary studies (comparing different MOIs) showed that at this MOI, there is a uniform distribution of the bacterial strain over the cell monolayer without causing any bacterial aggregates

Our results have shown the ability of *S. aureus* 8325.4 compared to *S. epidermidis* 1457 to adhere to bone marrow MSCs and differentiated osteoblasts, adipocytes and chondrocytes as shown in immunofluorescence adhesion assays in Figures 1, 2, 3, and 4. We found that FnBP plays a major role in this interaction as the adherent bacterial number was greatly reduced when using *S. aureus* ΔFnBP, a mutant lacking the expression of FnBP A/B, and then the percentage of adherent bacteria increase again, more than the wild type strain, when using bacterial strain ΔFnBP p4A that is ΔFnBP mutant strain supplemented with plasmid expressing intact FnBP type A as shown in Figures 1, 2, 3, and 4. The effect of ΔFnBP p4A may suggest that FnBP type A may play a greater role in bacterial adhesion.

FBS contains large number of extracellular matrix (ECM) proteins with high percentage of fibronectin and it has been found that ECM fibronectin plays an important role in the adhesion of *S. aureus* to several mammalian cell types, including MSCs, by serving as a bridge facilitating the interaction of bacteria FnBP with cells α5β1 integrin receptors (32). Our results showed that the presence of 10% FBS in infection medium results in high, specific adhesion between *S. aureus* WT and MSCs in contrast to the adhesion in the presence of serum free medium that results in high, non-specific and very low specific adhesion as shown in Fig. 5 (E) and (A). On the other hand, Fig.5 (C) and (G) showed greatly reduced bacterial adhesion in wells with *S. aureus* ΔFnBP in the presence of 10% FBS as a result of the absence of FnBP, while this was not the case in the presence of serum free medium with high, non-specific adhesion similar to that of the WT strain.

We have noticed that there is very low specific bacterial adhesion to MSCs in the presence of serum free medium, as stated above, and this may prove the release of cellular fibronectin by MSCs which is used as a bridge for the adhesion process. Also, we have found that the addition of plasma fibronectin at concentration of 400 μg/ml had very little effect on bacterial adhesion with high association control similar to serum free conditions as shown in Fig. 5 (I), (J), (K) and (L) of the results.

Lastly, the use of 10% decomplemented human serum results in very low bacterial number, as shown in Figure 5 (M), (N), (O) and (P) of the results, in both the adhesion wells and association control wells with weak immunofluorescence signal, and this may be attributed to the presence of precipitating anti staphylococcus antibodies in human serum and the possible role of Staphylococcus protein A (SPA) in inducing immune response.

The uptake of *S. aureus* by osteoblasts, and many other cell types necessitates the expression of FnBPs on the surface of the bacterium (6, 33, 34). These proteins belong to MSCRAMMs (microbial surface components recognising adhesive matrix molecules), which bind an array of extracellular matrix proteins such as collagen, elastin, fibrinogen, fibronectin, and bone sialo protein (35). The invasion of host cells by mutant bacterial strains lacking the two FnBPs, FnBPA and FnBPB were greatly reduced (33, 34). During the adhesion and invasion processes, *S. aureus* attaches to fibronectin via FnBPs expressed on their surface, and extracellular matrix fibronectin functions as a bridging molecule to the integrin α5β1 that serves as a “phagocytic” receptor leading to changes in the cytoskeleton of the cell and uptake of bacteria (9, 36).

Our results in the quantification association assay (Figures 6, 7 of results) showed that this specific binding protein plays a significant role in the binding of *S. aureus* to bmMSC. The two strains expressing this protein (WT, and ΔFnPB pA4) were able to adhere to and invade bmMSCs, while strains missing the binding protein failed to do that. ΔFnPB pA4, nevertheless, exhibited higher adherence followed by higher invasion results as paralleled to WT strain. Those results were consistent with other studies, as *S. aureus* was able to invade human Wharton’s jelly MSCs and adipose MSCs and that only small number of adherent bacteria were able to invade MSCs (5, 28).

The ability of *S. aureus* to interact with osteoblasts was extensively studied, mostly using cell lines. In this study we tested the ability of *S. aureus* to adhere and invade differentiated human osteoblasts (from bmMSCs) and we found that FnBP plays significant role in the interaction of *S. aureus* with osteoblasts (Fig. 8 and 9). The two strains expressing this protein (WT, and ΔFnPB pA4) was able to adhere to and invade osteoblasts, while strains missing the binding protein failed to do that. ΔFnPB pA4, nevertheless, exhibited higher adherence followed by nearly similar invasion results as paralleled to WT strain. These results are comparable to that of other studies (33, 37–40).

We also found that FnBP plays the same role in the interaction of *S. aureus* with differentiated adipocytes and chondrocytes (Figures 10, 11, 12 and 13 of results). The two strains expressing this protein (WT, and ΔFnPB pA4) was able to adhere to and invade these cells, while strains missing the binding protein failed to do that. Although, ΔFnPB pA4 does not shown significantly higher results than WT strain, as was the case with MSCs and to a less degree osteoblasts. A study by Hanses *et al*., 2011 confirmed the ability of *S. aureus* to invade adipocyte-like differentiated 3T3-L1 cells, but without studying the underlying mechanism for this invasion (41).

## Conclusion

Our results showed that S. aureus is able to interact with MSCs in the form of adhesion and invasion to the cells, and that this interaction is largely dependent on the expression of FnBP by S. aureus. We also showed that the same mechanism of interaction take place in case of S. aureus interaction with osteoblasts, adipocytes and chondrocytes that are directly differentiated from the same MSCs. Finally, we have found that the presence of 10% FBS in the infection medium is essential as it helps in achieving the best specific bacterial-cell association with the least background association.

## ACKNOWLEDGEMENTS

We are grateful to our sponsor, the HCED and the University of Bristol for their financial support. Thanks to Dr Andrew Edwards from University of Bath for providing us with the bacterial strains used in the experiments.

